# Live cell imaging of single neurotrophin receptor molecules on human neuron in Alzheimer’s disease

**DOI:** 10.1101/2020.02.17.953174

**Authors:** Klaudia Barabás, Julianna Kobolák, Soma Godó, Dávid Ernszt, Miklós Kecskés, Csaba Varga, József Kardos, Tibor Z. Jánosi, Takahiro Fujiwara, Akihiro Kusumi, Annamária Téglási, András Dinnyés, István M. Ábrahám

## Abstract

The changes in the receptor dynamics such as the surface movement of the receptor molecules on the plasma membrane are essential to receptor function. However, whether the receptor dynamics are affected by disease conditions is unknown. Neurotrophin receptors such as TrkA and p75^NTR^ play a critical role in neuronal survival and their functions are highly affected in Alzheimer’s disease (AD). Using live-cell single-molecule imaging of neurotrophin receptors we examined the surface trafficking of TrKA and p75^NTR^ molecules on human induced pluripotent stem cells (hiPSCs) derived live neurons from presenilin 1 (PSEN1) mutant AD patients and healthy subjects. Here we report that surface trafficking of p75^NTR^ molecules on neurites is faster than that of TrkA molecules in healthy controls. The surface dynamics of TrkA molecules were elevated in AD patients compared to healthy individuals. In contrast, the surface movement of p75^NTR^ was significantly smaller in AD patients compared to healthy individuals. Interestingly, amyloid beta_1-42_ (Aβ_1-42_) administration increased the surface trafficking of both TrkA and p75^NTR^ in healthy hiPSCs neurons. These findings provides the first evidence that the surface diffusion of TrkA and p75^NTR^ molecules are altered in patients suffering from AD. Our data also suggest that Aβ_1-42_ may responsible for the alteration of the surface movements of TrkA but not for p75^NTR^.

**One Sentence Summary:** The surface movements of neurotrophin receptors such as TrkA and p75^NTR^ are altered in neurons derived from patients suffering from familial Alzheimer’s disease.

## Introduction

Alzheimer’s disease (AD) is the most common type of dementia which is characterized by extracellular deposition of beta amyloid (Aβ) and intracellular neurofibrillary tangles *(1)*. The familial form of AD (fAD) is caused by genetic mutations (e.g. mutation in gene encoding presenilin (PSEN1 or PSEN2) *(2)*. Both sporadic (sAD) and fAD patient derived iPSC-based studies revealed increased Aβ_42_/Aβ_40_ ratios (*(3–6)*, elevated TAU hyperphosphorylation and increased GSK3B activity *(6–9)* recapitulating the major hallmarks of AD in an *in vitro* system, providing an effective platform to observe phenotypes relevant to AD in living human neurons.

Changes of cellular function in AD are due to the alteration in the signaling pathways such as low affinity binding p75 neurotrophin receptor (p75^NTR^) and tropomyosin receptor kinase A receptor (TrkA) related signaling. The unbalanced signaling through p75^NTR^ versus TrkA leads to neuronal loss in AD *(10, 11)*. When the ratio of p75^NTR^/TrkA is high, nerve growth factor can activate downstream signaling of p75^NTR^ and promote apoptosis *(12)*.

Signal transduction starts with receptor activation, which in turn can be precisely described by the changes of their surface trafficking in the plasma membrane *(13, 14)*. The neurotrophin receptors (e.g TrkA) have two distinct motility states on the membrane surface, characterized as mobile and immobile phases *(15, 16)*. The membrane recruitment of downstream intracellular signaling proteins such as MAPK occurs when TrkA is in the immobile phase *(15)*. Accordingly, the activation of neurotrophin receptors can be precisely measured by surface trafficking parameters of receptor molecules. However, how the surface trafficking of neurotrophin receptors in disease conditions is affected remains largely unknown.

Our understanding of AD pathogenesis is currently limited by difficulties in obtaining live neurons from patients *(17)*. Furthermore, the culture of primary human neuronal cells is particularly challenging because of their limited life span *(18)*. Human induced pluripotent stem cells (hiPSCs) based technology overcomes many of these limitations *(19)*.

Here we used neurons derived from iPSCs of PSEN1 mutant patients and non-dement control subjects. Using live cell single-molecule imaging of p75^NTR^ and TrkA we examined the surface trafficking parameters of neurotrophin receptor molecules in living human neurons and also observed whether the surface movement of the neurotrophin receptors are modified by Aβ_1-42_ treatment. Our results showed that the surface trafficking of p75^NTR^ molecules decreased in AD, while the surface movements of TrkA were increased. We also found that Aβ_1-42_ administration increased the diffusion of both TrkA and p75^NTR^ molecules in non-AD human neurons.

## Methods

### Chemicals and Reagents

All chemicals are from Sigma-Aldrich (St Louis, MO, USA) and the cell culture reagents and culture plates from Thermo Fisher Scientific (Waltham, MA, USA), unless specified otherwise.

### iPSC lines

Alzheimer’s disease (AD) patient derived iPSC line, BIOT-7183-PSEN1 (referred as BIOTi001-A in hPSCreg; *https://hpscreg.eu/*), bearing a mutation in PSEN1 gene (PSEN1 c.265G>C, p.V89L; characterized and published earlier *(20)* was labeled as fAD-1. IPSC line fAD-3 and fAD-4 were generated from two male siblings, bearing PSEN1 c.449T>C, p.L150P mutation, as published previously *(21)*. The patients were clinically diagnosed and characterized by the Institute of Genomic Medicine and Rare Disorders, Semmelweis University, Budapest (Hungary) or at the Danish Dementia Research Centre, Rigshospitalet, University of Copenhagen as described previously *(6, 20, 21)*. Non-demented volunteers (assessed by clinical evaluation) were used as controls (Ctrl-1; Ctrl-2; and Ctrl-5), from whom iPSC lines were established, characterized and maintained under identical conditions as the AD iPSC lines, as we published earlier (Ochalek et al. 2017). The hiPSC lines were maintained on Matrigel (BD Matrigel; Stem Cell Technologies) in mTESR1 (Stem Cell Technologies) culture media. The media was changed daily, and the cells were passaged every 5-7 days using Gentle Cell Dissociation Reagent, according to the manufacturer’s instructions.

### Neural induction of iPSCs

Neural progenitor cells (NPCs) were generated from each of the human iPSCs by dual inhibition of SMAD signaling pathway using LDN193189 and SB431542 *(22)*. Neural induction was initiated upon reaching approx. 90% confluence of iPSCs on Matrigel-coated dishes by addition of Neural Induction Medium (NIM) (1:1 (v/v) mixture of Dulbecco’s Modified Eagle’s/F12 and Neurobasal Medium, 1x N-2 Supplement, 1x B-27 Supplement, 1x Nonessential Amino Acids (NEAA), 2 mM L-Glutamine, 50 U/ml Penicillin/Streptomycin, 100 µM β-mercaptoethanol, 5 µg/ml insulin), which was supplemented with 5 ng/ml basic fibroblast growth factor (bFGF), 0.2 µM LDN193189 (Selleckchem) and 10 µM SB431542. The NIM medium was changed every day. At day 10 neural rosettes were picked manually and re-plated on poly-L-ornithine/laminin (POL/L; 0.003%/3 µg/cm^2^) coated dishes and expanded in Neural Maintenance Medium (NMM) (1:1 (v/v) mixture of Dulbecco’s Modified Eagle’s/F12 and Neurobasal Medium, 1x N-2 Supplement, 1x B-27 Supplement, 1x NEAA, 2 mM L-Glutamine, 50 U/ml Penicillin/Streptomycin), and supplemented with 10 ng/ml epidermal growth factor (EGF) and 10 ng/ml bFGF.

### Neural differentiation of NPCs

To generate human neurons, NPCs were plated on the POL/L coated dishes and cultured in Neural Differentiation Medium (NDM) (1:1 (v/v) mixture of Dulbecco’s Modified Eagle’s/F12 and Neurobasal-A medium, 1x N-2 Supplement, 1x B-27 Supplement, 1x NEAA, 2 mM L-Glutamine, 50 U/ml Penicillin/Streptomycin). For terminal differentiation into cortical neurons, the cells were plated on POL/L (0.002%/2 µg/cm^2^) at a seeding density of 30.000 cells/cm^2^ and 100.000 cells/cm^2^ for ELISA experiments with NMM medium. The medium was changed every 3-4 days during the terminal differentiation, until week 7. The efficiency of terminal differentiation was monitored by immunocytochemical staining for beta–III tubulin (TUBB3) and microtubule-associated protein 2 (MAP2) expression at week 7 (Fig. 1. A).

**Figure 1.**
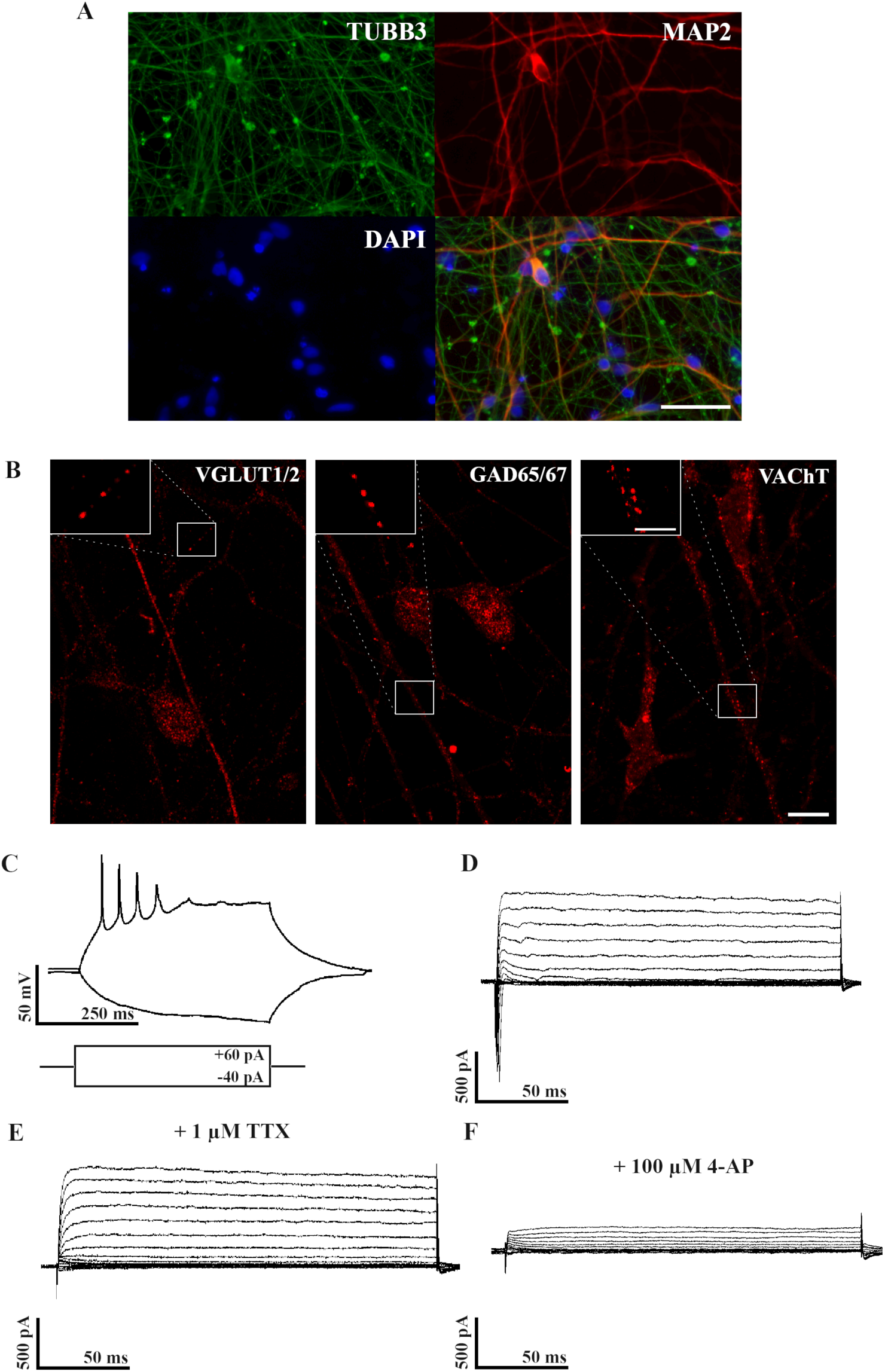
Neuronal phenotypes of differentiated hiPSCs. Immunofluorescence staining of microtubule-associated protein 2 (MAP2, red) and β–III tubulin (TUBB3, green) indicating that the hiPSC derived neuronal cultures show neuronal phenotypes by week 7 (A). Nuclei were labeld with DAPI (blue). Scale bar = 50 µm. Images depict VGLUT1/2, GAD65/67 and VAChT immunoreactivity in cytoplasm and neurites of 7 weeks old human iPSC derived neurons (B). Inserts demonstrate VGLUT1/2, GAD65/67 and VAChT immunoreactive dots on neurites of hiPSCs. Scale bar = 10µm, insert = 5 µm. Whole-cell patch-clamp recordings show that maturing neurons (7 weeks old) generate action potentials (C) and display Na^+^ and K^+^ currents (D) that can be blocked by TTX (E) or 4-AP (F), respectively.

### Preparation of Aβ_1-42_ peptide

Aβ_1-42_ was expressed recombinantly in *E. coli* and purified as it was described earlier *(23)*. Before the experiments, the monomeric lyophilized peptide was first dissolved in 10 mM NaOH on ice, then it was diluted into appropriate amount of TBS buffer resulting a final solution of 300 µM peptide concentration and a pH of 7.4. The solution was aged for 4 days at room temperature. Before administration, the solution was further diluted in NMM medium to the required concentration and was spun down in a tabletop centrifuge at 15,000 g for 2 minutes to remove larger aggregates.

### Measurement of Aβ_1-40_ and Aβ_1-42_ by ELISA

Conditioned media was collected after 4 days of culture (without media change) at week 7 from one well of a six-well plate. To prevent protein degradation, 4-(2-Aminomethyl) benzenesulfonyl fluoride hydrochloride (AEBSF) was added to the media. Extracellular Aβ_1-40_ and Aβ_1-42_ levels were measured using Human β-Amyloid (1-40) ELISA Kit and Human β-Amyloid (1-42) ELISA Kit (Wako), according to the manufacturer’s instructions. The secreted Aβ levels determined in pM were normalized to the total protein content (µg) of cell lysate. Total protein concentration was determined using a Pierce BCA Protein Assay Kit, where cells were lysed with RIPA Lysis and Extraction Buffer supplemented with Halt™ Protease and Phosphatase Inhibitor Cocktail and Pierce™ Universal Nuclease for Cell Lysis. Lysed samples were sonicated and centrifugated at 13000 rpm to collect the supernatants. The signal was detected with Varioskan Flash Multimode Reader (Thermo Fisher Scientific). As all fAD cell lines (fAD-1, fAD-3, and fAD-4; bearing PSEN1 mutations) showed significant difference (p<0.01) from controls (Ctrl-1, Ctrl-2, and Ctrl-5) and they were not significantly different from each other we show the results of pooled data.

### Immunocytochemistry and image analysis

Single-label immunofluorescence was performed to detect neuronal phenotypes (MAP2, TUBB3, VGLUT1/2, GAD65/67, VAChT) and examine the expression of TrkA or p75^NTR^ in terminally differentiated iPSCs derived neurons originated from healthy individuals and fAD patients. Cells were fixed with 4% PFA (pH 7.4) for 20 minutes, then were pretreated with TRIS containing 1-3% bovine serum albumin (Sigma) and 0.1-0.3% Triton-X depending on the detectable target protein. Cells then were incubated in the appropriate primary antibodies (Table S1) overnight at 4°C or 2 hours at room temperature (RT). To detect the signal, cells were incubated 60-120 minutes at RT with the appropriate secondary antibodies (Table S1). Cell nuclei were visualized using Vectashield Mounting Medium with DAPI (1.5 µg/mL; Vector Laboratories). MAP2, TUBB3, VGLUT1/2, GAD65/67, VAChT, TrkA and p75^NTR^ positive cells were analyzed under fluorescent microscope equipped with 3D imaging module (Axio Imager system with ApoTome; Carl Zeiss MicroImaging GmbH) controlled by AxioVision 4.8.1 software (Carl Zeiss) or under confocal laser scanning microscope (Zeiss LSM710) (CLSM). Helium neon laser with 633-nm wavelength was used to excite Alexa 647. The diameter of the pinhole aperture was set to gain an optical thickness of 1 µm. Optical density was measured on neurites for estimation of quantitative differences in TrkA and p75^NTR^ immunoreactivity between experimental groups. Ten control and ten AD confocal images were analyzed at 60x magnification with an investigator blind to the experimental groupings. Using the FIJI package of the ImageJ software the OD value was obtained by averaging OD values of each selected region of interest (ROI=108 µm^2^) after background subtraction and grey scale threshold determination. OD values of control and PSEN1 mutant neurites were paired in case of both TrkA and p75^NTR^ and the ratio between the OD values of TrkA and p75^NTR^ (TrkA/p75^NTR^) was calculated.

### Electrophysiology

Electrophysiological properties of terminally differentiated human iPSCs derived neurons were tested using whole-cell patch-clamp recording. Patch pipettes were pulled from borosilicate glass capillaries with filament (1.5 mm outer diameter and 1.1 inner diameter; Sutter Instruments) with a resistance of 2-3 MΩ. The pipette recording solution contained (in mM) 10 KCl, 130 K-gluconate, 1.8 NaCl, 0.2 EGTA, 10 HEPES, 2 Na-ATP, pH 7.3 adjusted with KOH. All recordings were performed at 32 °C with the chamber perfused with oxygenated ACSF containing (in mM) 2.5 KCl, 10 Glucose, 126 NaCl, 1.25 NaH_2_PO_4_ 2 MgCl_2_, 2 CaCl_2_ 26 NaHCO_3_. Whole-cell recordings were made with Axopatch 700B amplifier (Molecular Devices) using an upright microscope (Nikon Eclipse FN1) equipped with differential interference contrast optics (DIC). Cells with access resistance below 20 MΩ were used for analysis. Signals were low-pass filtered at 5 kHz and digitized at 20 kHz (Digidata 1550B, Molecular Devices). Acquisition and subsequent analysis of the acquired data were performed using Clampex9 and Clampfit software (Axon Instruments). Traces were plotted using Origin8 software (MicroCal Software, Northampton, MA).

### TrkA and p75^NTR^ labelling of live neurons and Aβ_1-42_ treatment

Live cell immunocytochemistry was performed to detect TrkA and p75^NTR^ molecules on living neurons. Terminally differentiated human iPSCs derived neurons were grown on Poly-L-ornithine and Laminin (POL/L; 0.002%/3 µg/cm^2^) coated 35 mm glass-bottom dish (MatTek Corporation) in NMM medium. Neurons were incubated with Atto-633 labeled anti-TrkA antibody (anti-TrkA (extracellular)-Atto-633, 1:100, Alomone Labs) or Atto-488 labeled anti-p75^NTR^ antibody (anti-p75^NTR^ (extracellular)-Atto-488,1:100, Alomone Labs) for 6 minutes at 37°C.

The specificity of the TrkA and p75^NTR^ antibodies have been validated by preadsorption of the primary antibody with its corresponding fusion protein and the specificity of TrkA has also been tested in TrkA expressing Chinese hamster ovary (CHO) cells. TrkA expressing CHO cells showed labeling with anti-TrkA Atto-633 antibody, while native (non-TrkA expressing) CHO cells showed a complete absence of the signal.

For the Aβ_1-42_ treatment hiPSC neurons from non-demented volunteers were labelled with anti-TrkA and anti-p75^NTR^ antibody as described above, then the washing medium was replaced by NMM medium containing 1 µM aged Aβ_1-42_. Single-molecule imaging was commenced when Aβ_1-42_ was administered and was carried out for 60 minutes.

### Single-molecule imaging of TrkA and p75^NTR^ molecules

Single-molecule imaging of labeled TrkA and p75^NTR^ molecules was carried out on an Olympus IX81 fiber total internal reflection fluorescence microscope equipped with ZDC (zero drift compensation) stage control and Plan Apochromat objective (100x, NA 1.45, Olympus) and a humidified chamber (Supertech) heated up to 37°C containing 5% CO_2_. Diode lasers (Olympus) was used to excite ATTO-488 at 491 nm and ATTO-633 at 640 nm wavelength and emission was detected above 510 nm and at 650-670 nm emission wavelength range, respectively. EM CCD camera (Hamamatsu 9100-13) and imaging software (Olympus Excellence Pro) was applied for image acquisition. Image sequences were recorded for 60 minutes in case of each sample. 20-50 images were recorded for 10 seconds long sampling interval with 33 ms acquisition time. Critical angle of the laser beam was adjusted to set up the penetration depth to 100 nm. The number of fluorophores per analyzed spots was determined with single-step photobleaching. (Supplementary Fig. 3.).

Single-molecule tracking was completed in a custom-made software written in C++. The center of each fluorescent signal was localized by two-dimensional Gaussian fitting. A minimum step size linking algorithm connected the localized dots in the image series and generated trajectories. Trajectories were examined one by one and artefacts or tracks shorter than 15 frames were excluded from the subsequent analysis.

In order to identify the examined neurotrophin receptor molecules expressed by neuronal phenotypes, correlated live cell single-molecule imaging and fixed cell immunocytochemistry were performed. The x/y coordinates of in vivo labelled TrkA and p75^NTR^ positive neurites were identified by means of the imaging software and x/y memory of the motorized microscope stage after the single-molecule experiments. Cells then were fixed with 4% PFA and beta-III tubulin immunocytochemistry was performed in the same manner as detailed above. The image sequences of the in vivo labelled receptors and beta-III tubulin images were merged using FIJI software and observed whether the TrkA and p75^NTR^ molecules were moving along beta-III tubulin immunopositive neurites (Supplementary Movie 3.). Our findings indicate that each receptor molecules of our single-molecule imaging analysis are expressed by neurons.

The viability of the cells was tested with a LIVE/DEAD viability/Cytotoxicity Assay Kit (Thermo Fisher Scientific) at the end of experiments according to the manufacturer’s instructions. The results demonstrated that cells retained their plasma membrane integrity until the end of the experiments (Supplementary Fig. 1.).

### Diffusion coefficient (D) calculation with the maximum likelihood method

Maximum-Likelihood estimation *(24)* was applied to obtain the corresponding diffusion coefficient for each trajectory. Δ represents the observed displacements 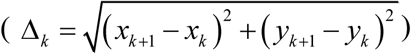, arranged in an *N*-component column vector where the total number of frames is equal to *N*+1. *x*_n_ and *y*_n_ are the coordinates of the signal’s centre on the *n*th frame.

Σ is the *N* x *N* covariance matrix, defined by the following equation *(25)*:

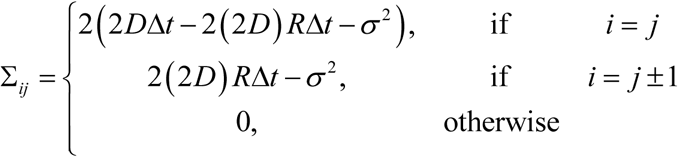

where D is the diffusion coefficient, Δ*t* is the frame integration time, *σ* is the static localization noise, *R* summarizes the motion blur effect and in our case *R*=1/6 as a consequence of the uniform illumination.

The likelihood function was defined by the following function:

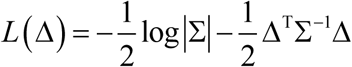

The *D* and *σ* which provides the maximal *L*(Δ) value is the estimated diffusion coefficient and static localization noise respectively. This global optimization can be carried out directly. However, in this case it is necessary to calculate the determinant and the inverse of covariance matrix at each step of the optimization method. In this particular case (*N* can be more than 200 this computational inconvenience becomes severe difficulty. Fortunately, an approximation does exist *(26)* which based on the theory of circulant matrices and it is applicable *(25)*: if the total number of frames is sufficiently high. The global optimization was performed by a numerical method implemented in MATLAB. The goodness of optimization was judged by the evaluation of the static localization noise. An optimization was considered to be inaccurate and was excluded from further analysis when the estimated static localization noise was out of ±90% range of the group’s mean.

### Statistical analysis

Values are expressed as mean ± SEM. Dunnett’s method was performed to compare the individual groups to controls in ELISA experiments. Kolmogorov-Smirnov test was used to compare the distributions of D values and Mann-Whitney U test was applied to compare OD values. Statistical differences were considered significant at *p<0.05; **p<0.01; ***p<0.001. All results were analyzed using the GraphPad Prism 5 software and Statistica 13.3 for Windows (TIBCO).

## Results

### Characterization of the hiPSC derived neurons

We tested whether the iPSC derived neurons show neuronal characteristics comparable with human mature neurons. First, we stained the fixed terminally differentiated neuronal cultures with tubulin, beta 3 class III (TUBB3) and microtubule-associated protein 2 (MAP2). All the cell lines showed neuron-like morphology and positive staining for the neuronal markers (Fig.1. A), representing the neuronal differentiation of the iPSC derived cell cultures.

We also identified the neurochemical specificity of our 7-weeks old hiPSC derived neurons by immunolabeling them for glutamatergic (VGLUT1/2), GABAergic (GAD65/67) and cholinergic (VAChT) markers. We found that our hiPSC neurons represent a phenotypically mixed population including glutamatergic, GABAergic and cholinergic neurons (Fig.1. B).

Next, we examined whether our neuronal differentiation protocol resulted in cells with electrophysiological properties similar to mature neurons. Using whole-cell patch-clamp technique, we recorded action potentials and voltage-gated Na^+^ and K^+^ currents from seven weeks old neuronally differentiated cells. As evidence for maturation, cells exhibited either single or repetitive action potentials in response to positive current injection (Fig.1. C). Furthermore, a series of 10 mV depolarization voltage steps from −90 mV resulted in the opening of Na^+^ and K^+^ channels indicated the fast inward and slow outward component, respectively (Fig.1. D). TTX blocked the inward, while 4-AP the outward currents illustrating that the currents were a result of the activation of voltage-gated Na+ and K+ channels (Fig.1. E, F).

### Aβ production by healthy and PSEN1 mutant hiPSC neuron

We determined the Aβ production of the 7 weeks old neuronal cultures by measuring the secreted Aβ_1-40_ and Aβ_1-42_ levels in the conditioned media of differentiated neural cells by ELISA. We detected significantly elevated Aβ_1-40_ and Aβ_1-42_ levels in the PSEN1 mutant cultures, and significantly increased Aβ_1-42_/Aβ_1-40_ ratios (p<0.01) compared to the control hiPSC neuron from non-dement volunteers (Supplementary Fig.2.). Based on our results all fAD cell lines (fAD-1, fAD-3, and fAD-4; bearing PSEN1 mutations) showed significant difference (p<0.01) from controls.

### Expression and surface movement of TrkA and p75^NTR^ in healthy and PSEN1 mutant hiPSC neurons

Our immunohistochemical experiments demonstrated that TrkA and p75^NTR^ are expressed in neurites and somas (Fig. 2. A). Quantitative image analysis performed on fixed cultures revealed that the intensity of TrkA and p75^NTR^ immunoreactivity is increased in PSEN1 (Fig. 2. B) and the ratio of p75^NTR^ /TrkA elevated in PSEN1 mutant neurons (Fig. 2. C).

**Figure 2.**
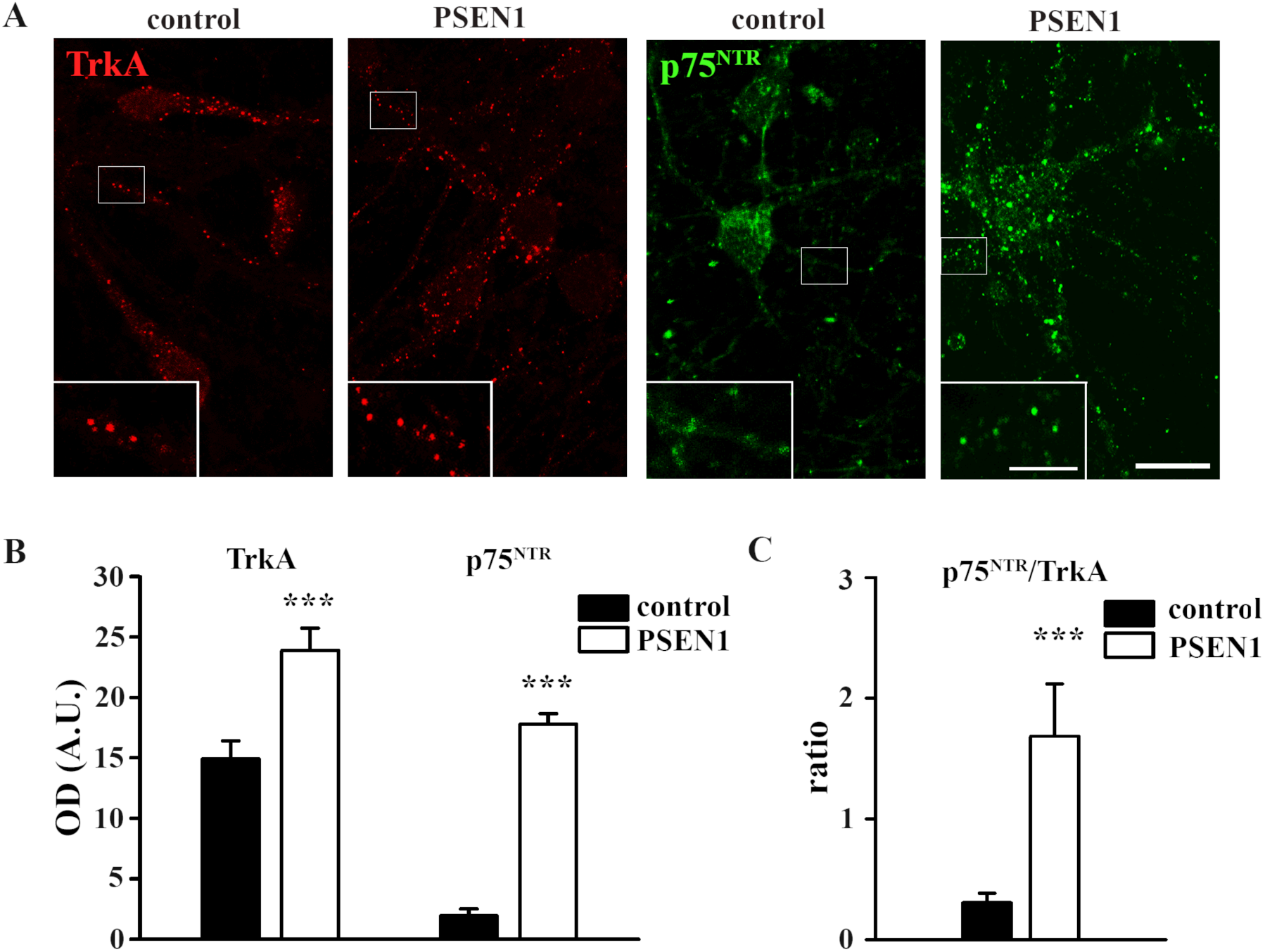
Characterization of TrkA and p75^NTR^ expression in healthy and PSEN1 mutant neurons. Images show TrkA (red) and p75^NTR^ (green) expression in the cytoplasm and neurites of 7 weeks old healthy and PSEN1 mutant neurons (A). Histograms demonstrate TrkA (A), p75^NTR^ (B) expression in PSEN1 mutant neurons compared to controls and their ratio (C) (***p<0.001). Data are expressed in optical densities (pixel/area) ± SEM. Scale bar = 20 µm, scale bar insert = 5 µm.

The fluorescence intensity versus time function showed one-step photobleaching representing single Atto-488 or Atto-633 fluorophore for p75^NTR^ and TrkA, respectively. The peak intensity of the intensity spot frequency histograms of both p75^NTR^ and TrkA were similar to that of the step sizes for photobleaching. These results suggest that most of the spots represented single fluorophores and single receptors (Supplementary Fig. 3). Based on the mean square displacement functions of TrkA and p75^NTR^, besides some immobile receptors, mainly two types of receptor movements were found (Fig.3. A, B). Brownian diffusion, when receptors are moving freely between barriers, and confined motion when receptors are restricted to a small area. The cumulative probability functions of D values of TrkA and p75^NTR^ on healthy neurites indicate that p75^NTRs^ are moving significantly faster on neurites than TrkAs on control non-demented neurons (Fig. 3. C).

**Figure 3.**
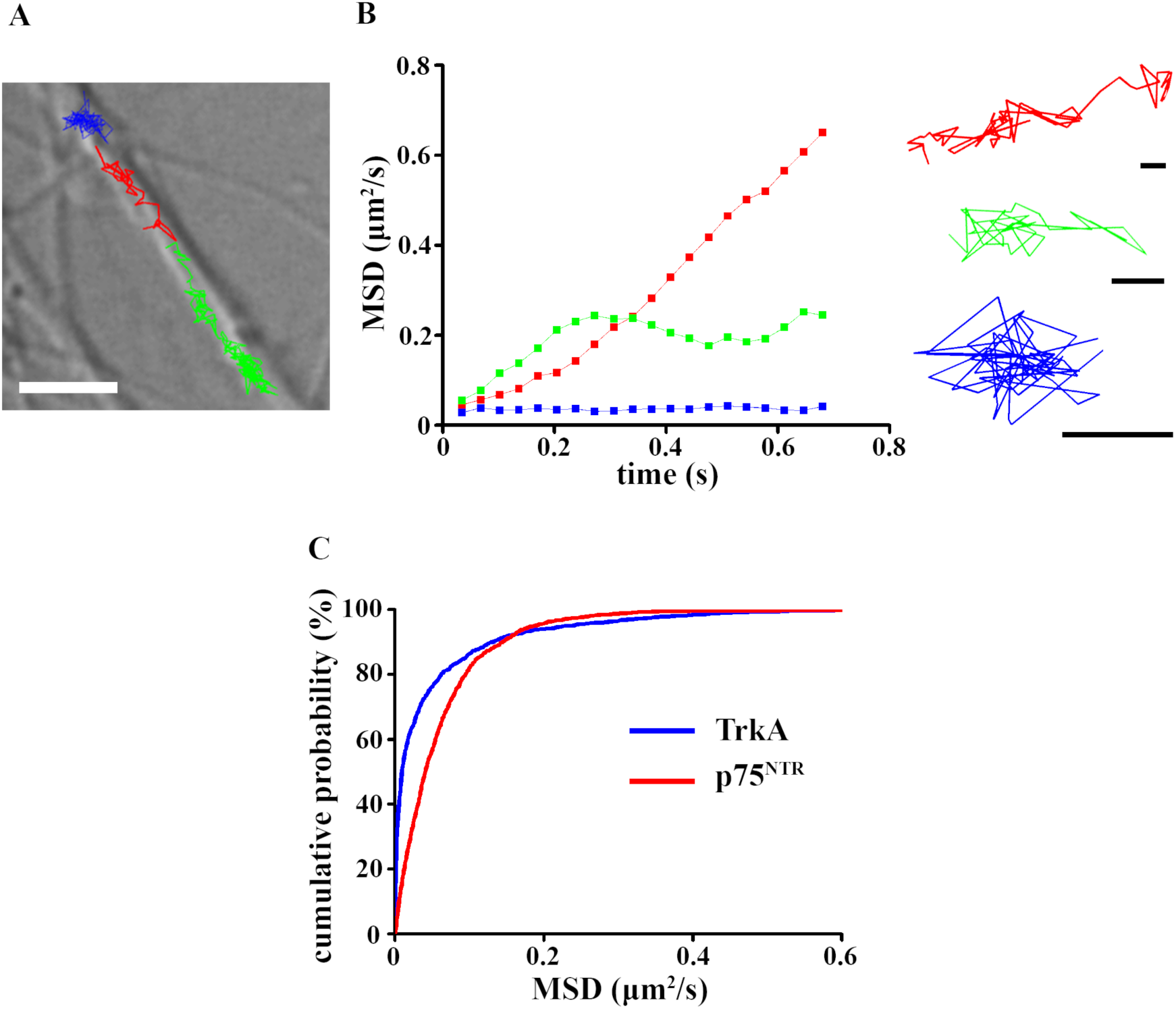
Single-molecule imaging of TrkA and p75^NTR^ in healthy neurons. Representative trajectories of TrkA molecules on neurites (A). Scale bar: 5 µm. The mean square displacement (MSD) functions represent TrkA molecules with different diffusion modes (B). Scale bars: 0.5 µm. The cumulative probability functions of D values of TrkA and p75^NTR^ on healthy neurites are shown (number of trajectories= 1556-1872) (C).

The distribution of D values showed that the surface movement of TrkA molecules increased in PSEN1 mutant neurites compared to non-demented controls (Fig.4. A). In contrast, the surface trafficking of p75^NTR^ receptor molecules decreased in PSEN1 mutant neurites compared to controls (Fig.4. B).

**Figure 4.**
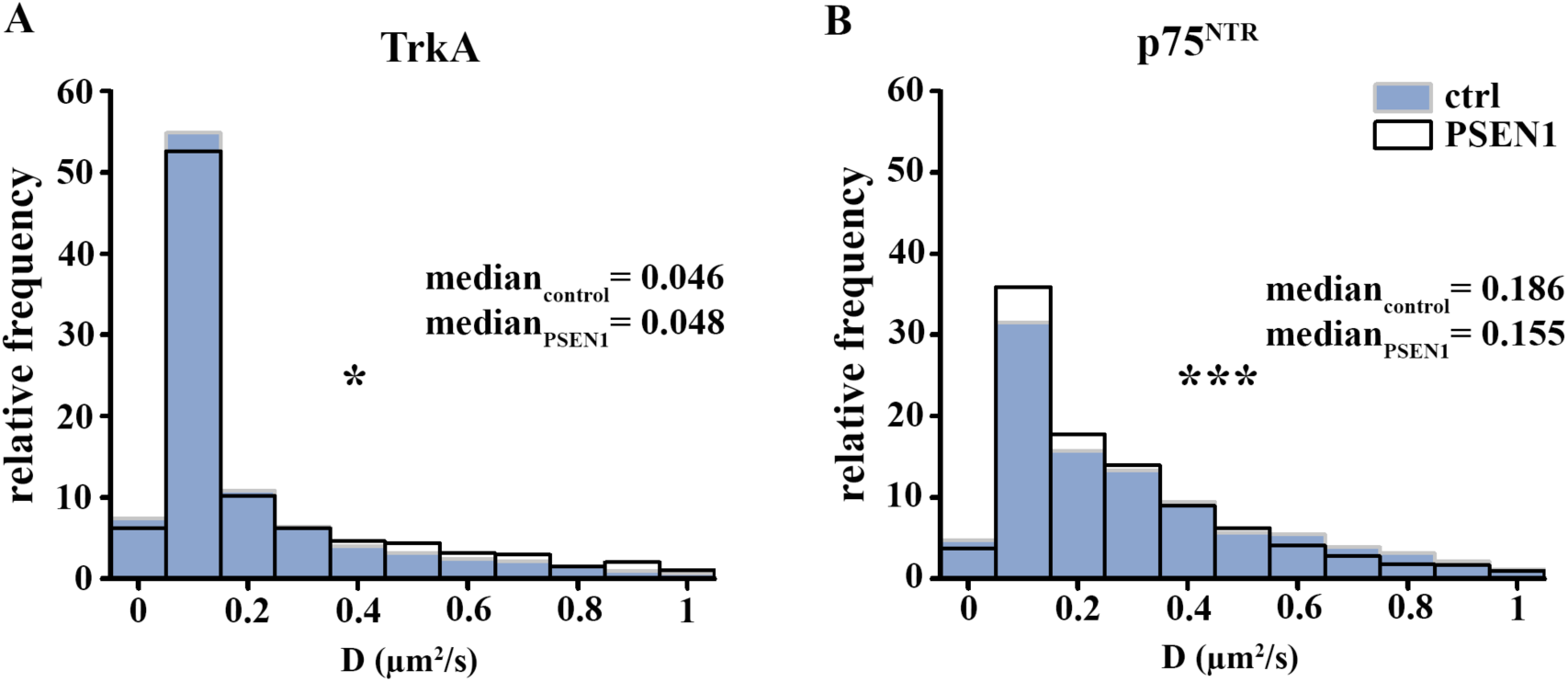
Distribution of the diffusion coefficients of TrkA and p75^NTR^ in healthy and PSEN1 mutant neurons. Histograms show the relative frequency of different diffusion intervals in the range of 0-1 µm^2^/s for TrkA (A) and p75^NTR^ (B) in non-dement, control (blue bars) and PSEN1 mutant neurites (white bars). (TrkA: p=0.0156; number of trajectories=1556-1378, p75^NTR^: p=0.0002; number of trajectories=1872-2148.)

D values from control non-demented (Ctrl-1, Ctrl-2, and Ctrl-5) and fAD cell lines (fAD-1, fAD-3, and fAD-4; bearing PSEN1 mutations) were pooled and analyzed together.

### Effect of Aβ_1-42_ on surface movement of TrkA and p75^NTR^ in healthy neurons

In ordert to detect the effect of Aβ_1-42_ on surface movement of TrkA and p75^NTR^ single molecua imaging experiments were performed on the control cell lines from non-demented volunteers. Application of 1µM Aβ_1-42_ significantly increased the surface trafficking of both TrkA and p75^NTR^ in non-mutant neurites (Fig.5. A, B).

**Figure 5.**
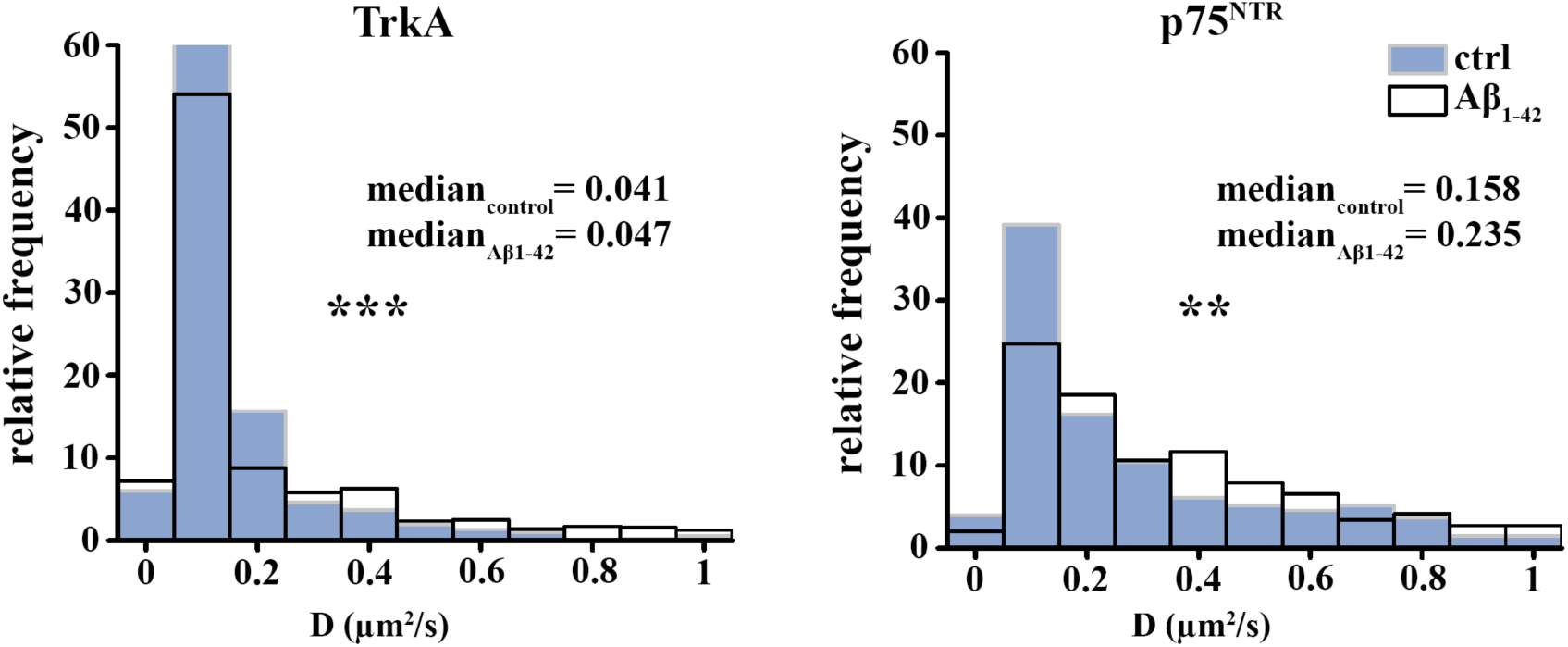
Distribution of the diffusion coefficients of TrkA and p75^NTR^ in healthy and Aβ_1-42_-treated healthy neurons. Histograms show the relative frequency of different diffusion intervals in the range of 0-1 µm^2^/s for TrkA (A) and p75^NTR^ (B) in non-dement, control (blue bars) and 1 µM Aβ_1-42_-treated neuronal fibres (white bars). TrkA: p=0.0000018; number of trajectories=517-637, p75^NTR^: p=0.0002; number of trajectories=258-491.

## Discussion

Our results show that the surface trafficking of TrkA increases, while the trafficking of p75^NTR^ decreases in hiPSC neurons derived from PSEN1 mutant patients. We also demonstrate that the effect of acute Aβ_1-42_ administration to healthy hiPSCs resembles to the changes seen in the surface trafficking of TrkA but does not mimic the changes observed in the movement of p75^NTR^ in PSEN1 mutant neurons.

First, we have characterized the neuronal phenotype of the hiPSCs. Our results demonstrated that 7 weeks old hiPSCs express neuronal markers such as TUBB3 and MAP2 and show electrophysiological characteristics of mature neurons. In addition, we tested their neuronal phenotype and revealed that hiPSCs form a mixed population of glutamatergic, GABAergic and cholinergic neurons. We have also validated the fAD patient’s iPSC-based cellular pathology model used in this study. Our results showed that PSEN1 mutant hiPSC neurons produce more Aβ_1-40_ and Aβ_1-42_ and Aβ_1-42_/Aβ_1-40_ ratio increased compared to control cells. Previously we have confirmed that hyperphosphorylation of TAU protein and an increased level of active glycogen synthase kinase 3 beta (GSK3B), a physiological kinase of TAU, is detected in PSEN1 mutant patient groups *(6)*. In agreement with previous results *(6)*, our findings demonstrated that γ-secretase inhibitor DAPT treatment on healthy and PSEN1 mutant iPSC-derived neurons resulted in reduced endogenous amyloid levels and intracellular accumulation of AβPP-C-terminal fragment *(6)*.

In the current study, we investigated the surface trafficking of two neurotrophin receptors, TrkA and p75^NTR^ on healthy and PSEN1 mutant iPSC-derived neurons. The TrkA and p75^NTR^ play a critical role in the progress of AD by shifting the balance from TrkA-mediated survival to p75^NTR^–mediated pro-apoptotic signaling. Although the expression levels of TrkA and p75^NTR^ are differently affected depending on the examined brain area and the progression stage of AD *(11)*, their ratio is essential in determining the functional outcome. Our data show that both TrkA and p75^NTR^ levels are elevated in PSEN1 mutant neurites but the ratio of p75^NTR^/TrkA is higher in PSEN1 mutant neurons compared to neurons derived from non-dement volunteers suggesting that the balance is shifted toward the p75^NTR^ activated intracellular signaling in neurons from PSEN1 mutant patients.

The surface movements of the receptors indicate their activation state *(13, 14)*. Although the expression pattern of TrkA and p75^NTR^, their interaction and the stimulated signaling pathways leading to AD progression is relatively well known *(27, 28)*, their surface movements in AD remained unexplored. Here we showed that both TrkA and p75^NTR^ exhibit various diffusion modes in healthy hiPSCs and that p75^NTR^ moves faster along the neurites than TrkA. TrkA, however, moves faster in PSEN1 mutant neurons compared to controls. Contrarily, the movement of p75^NTR^ becomes slower in PSEN1 mutant hiPSCs. As described earlier TrkA is immobilized upon ligand binding, which is related to the start of signal transduction *(15)*. p75^NTR^ has also been shown to slow down and start signalization upon ligand binding *(16)*. Based on these results our data indicate that TrkA might be less, while p75^NTR^ might be more active in PSEN1 mutant neurons. However, the actual changes of their intracellular downstream action require further investigations and is beyond the scope of this study.

Since AD is associated with the formation of extracellular amyloid plaques *(29)* and PSEN1 mutation induce an excessive production of Aβ_1-42_ *(30)*, we examined whether the alteration in the lateral diffusion of TrkA and p75^NTR^ observed in our AD model is due to the increased Aβ_1-42_ levels. We treated hiPSC neurons derived from healthy individuals with 1 µM Aβ_1-42_. Our unpublished result showed that application of 1 µM Aβ_1-4_ resulted in significant hiPSC neuronal loss after 24 hours of application. The acute treatment of Aβ_1-42_ increased the diffusion coefficients of both neurotrophin receptors. These results suggest that the accumulation of Aβ_1-42_ might be involved in the alteration of the diffusion properties of TrkA seen in PSEN1 mutant neurons, while the change of diffusion parameters of p75^NTR^ in PSEN1 mutant cells is not mimicked by Aβ_1-42_.

In summary, our data provide first evidence that the surface trafficking of TrkA and p75^NTR^ is altered in PSEN1 mutant hiPSC neurons. Our results also suggest that altered surface trafficking of p75^NTR^ can not be explained by the presence of Aβ_1-42_ in fAD per se. Finally, these results draw attention to the significance of investigation of receptor dynamics in disease conditions.

## Acknowledgments

This work was supported by the Hungarian Brain Research Program (KTIA_NAP_13-2014-0001, 20017-1.2.1-NKP-2017-00002), the Hungarian Scientific Research Fund (OTKA; 112807), and the European Union and was co-financed by the European Social Fund under the following grants: EFOP-3.6.1.-16-2016-00004 (Comprehensive Development for Implementing Smart Specialization Strategies at the University of Pécs), EFOP 3.6.2-16-2017-00008 (The Role of Neuro-inflammation in Neurodegeneration: From Molecules to Clinics), the Higher Education Institutional Excellence Program of the Ministry for Innovation and Technology in Hungary, within the framework of the 5. thematic program of the University of Pécs and ÚNKP-18-3-III (New National Excellence Program of the Ministry of Human Capacities).

## Supplementary figures

**Supplementary figure 1.**
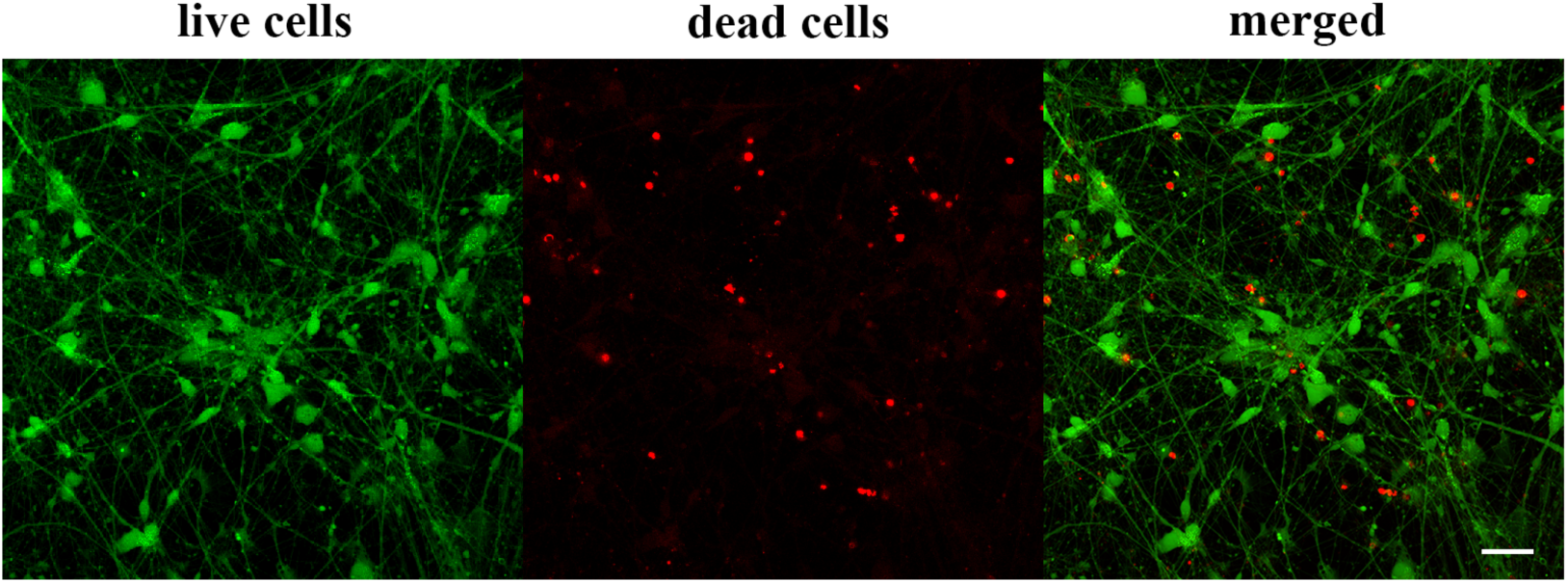
Viability of hiPSCs after measurements. Representative image shows that hiPSC neurons are alive after an hour measurement (A) Live cells are shown in green, dead cell population is labeled with red (green = calcein AM, red = ethidium homodimer-1).

**Supplementary figure 2.**
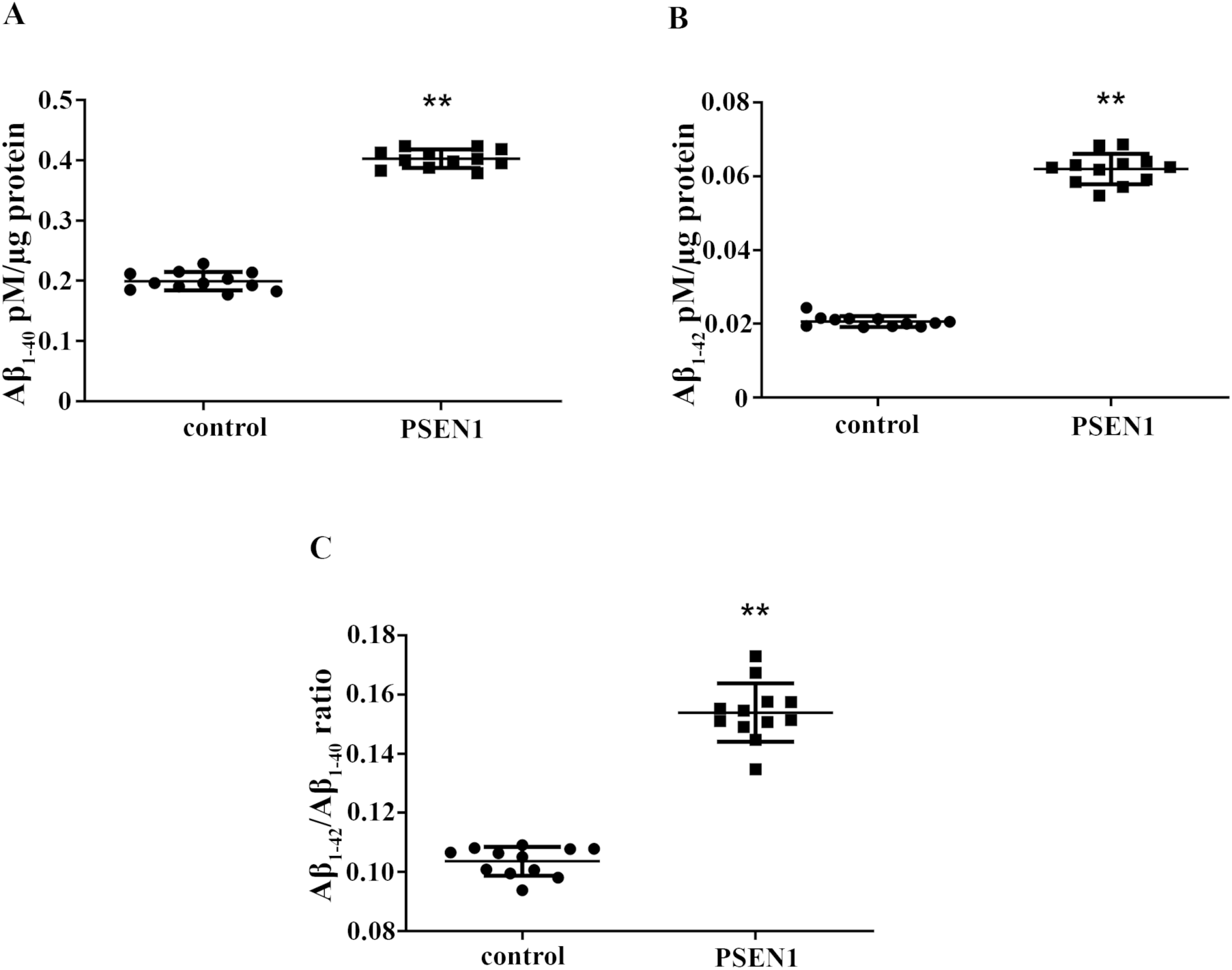
Levels of Aβ_1-40_ and Aβ_1-42_ in healthy and PSEN1 mutant neurons. Histograms show the levels of Aβ_1-40_, (A) Aβ_1-42_ (B) and the ratio of Aβ1-_42_/Aβ1-_40_ (C) in control and PSEN1 mutant neurons. The extracellular Aβ levels determined in pM were normalized to total protein content (in µg) of cell lysates. Data represent mean ± SEM (n=12). (**p<0.01).

**Supplementary figure 3.**
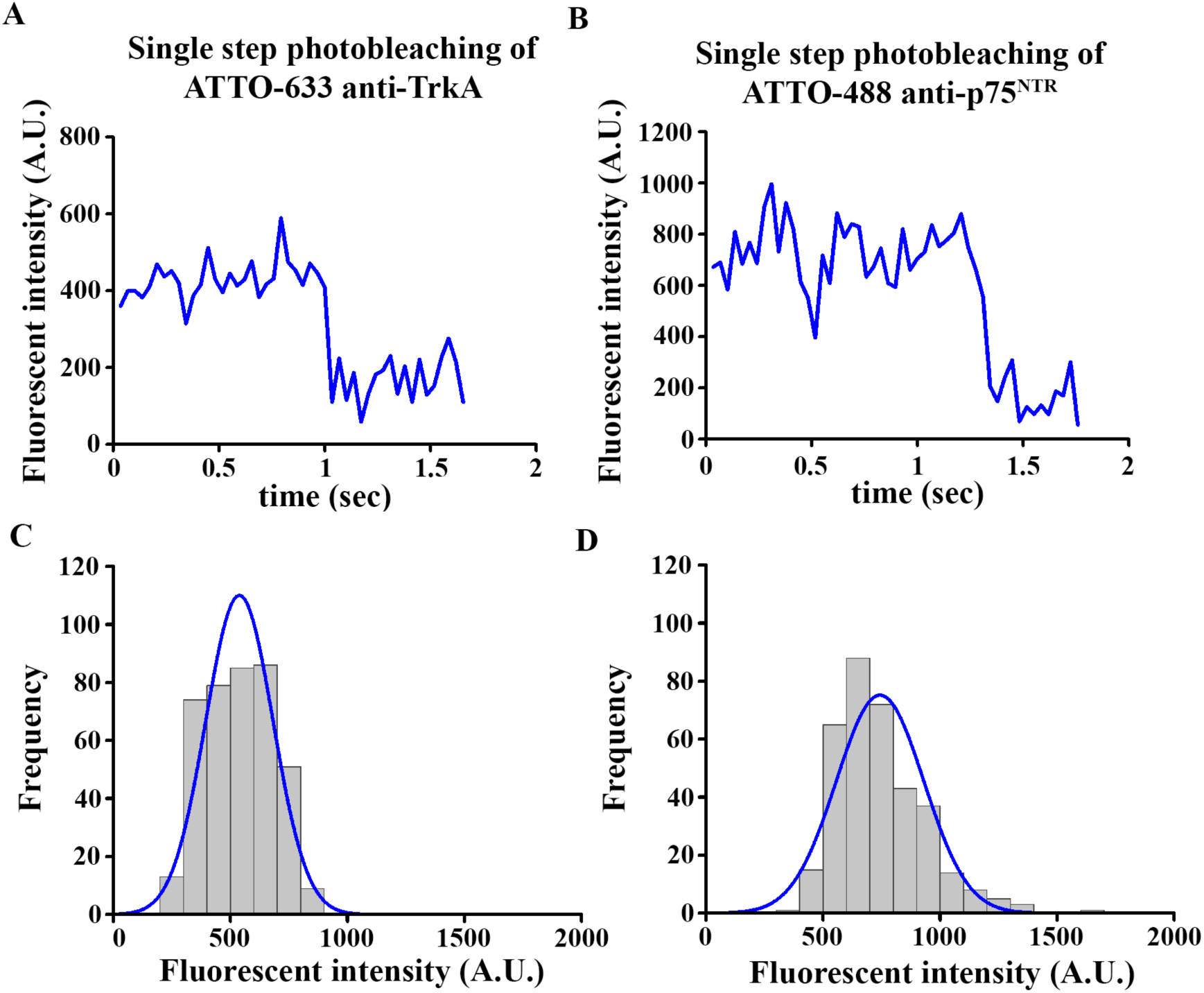
Single-step photobleaching of Atto-633 anti-TrkA and Atto-488 anti-p75^NTR^. Intensity profiles of a single Atto-633-labeled TrkA (A) and Atto-488-labeled p75^NTR^ (B) signal. Histograms show the intensity value of every spot for TrkA (C) and p75^NTR^ (D) in a recording superimposed with a single-fitted Gaussian curve (blue line).

**Supplementary table 1.**
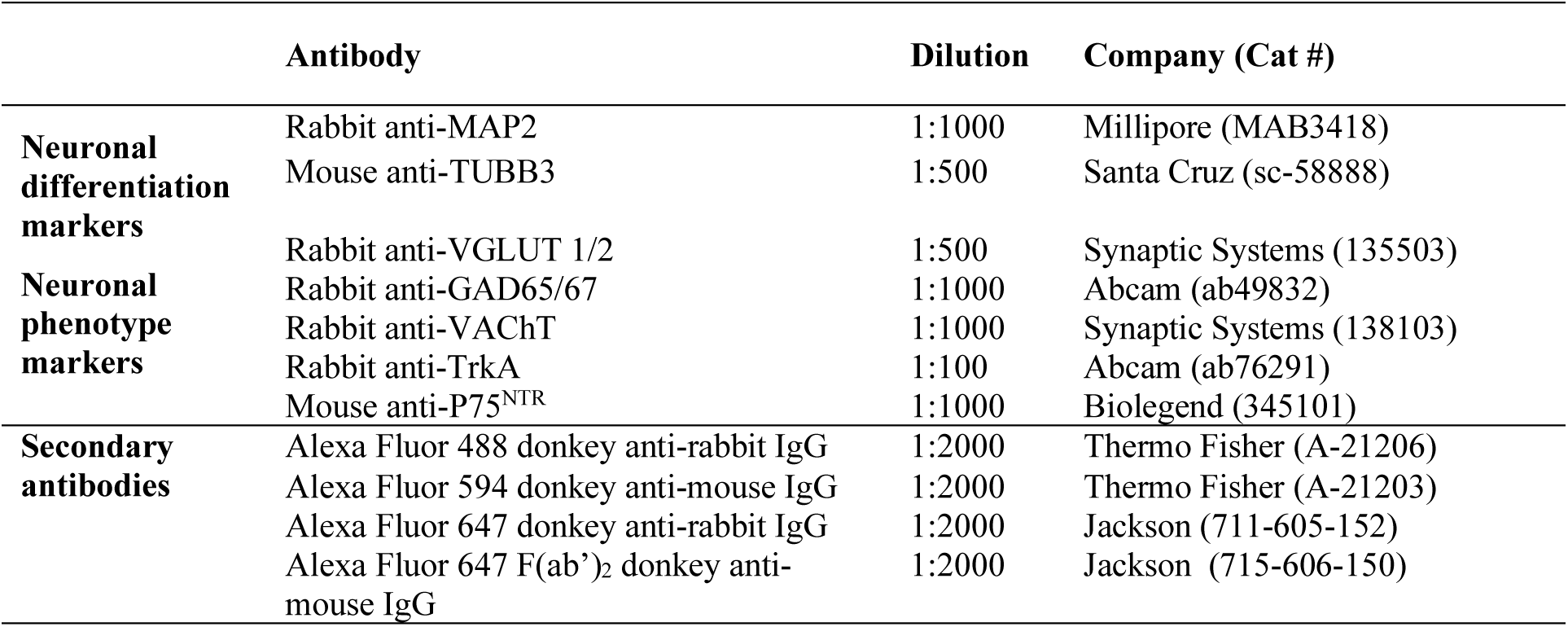
The table shows antibodies used for immunocytochemistry in fixed cells.

## Supplementary movies

**Movie 1. Surface movements of TrkA in healthy hiPSCs.** Single molecules of Atto-633-labeled TrkA are moving along a neurite in live hiPSC neuron from non-demented individuals. The recording was made in TIRF mode with a 34-ms acquisition time and is displayed at 29 fps. Scale bar: 1µm

**Movie 2. Surface movements of p75**^**NTR**^ **in healthy hiPSC neurons**

Single molecules of Atto-488-labeled p75^NTR^ are moving along a neurite in live hiPSC neurons from non-demented individuals. The recording was made in TIRF mode with a 34-ms acquisition time and is displayed at 29 fps. Scale bar: 1µm

**Movie 3. Correlated live cell single-molecule imaging and immunocytochemistry on fixed cell.** Representative video shows that Atto-488-labeled p75^NTR^ molecules (green) are moving along beta-III tubulin immunopositive neurites (red). The recording was made in TIRF mode with a 34-ms acquisition time and is displayed at 29 fps. Scale bar: 1µm

